# Water stress and CO_2_ concentration interactions affect carbon isotope signatures of leaf and phloem organic matter

**DOI:** 10.1101/252213

**Authors:** Yonge Zhang, Xinxiao Yu, Lihua Chen, Guodong Jia, Hanzhi Li

## Abstract

Investigation of δ^13^C of leaf and twig phloem water-soluble organic material (WSOM) is a promising approach for analysis of the effects of environmental factors on plant performance. In this study, orthogonal treatments of three CO_2_ concentrations (C_a_) × five soil water contents (SWC) were conducted using *Platycladus orientalis* saplings to investigate the interaction of water stress and CO_2_ concentration on δ^13^C of leaf and twig phloem WSOM. Under the lowest SWC, the δ^13^C of leaf and twig phloem WSOM had the most positive values at any C_a_ and their values decreased as C_a_ increased. However, at improved soil water conditions, the greatest values of δ^13^C of leaf and twig phloem WSOM were mostly observed at C_600_. In addition, a more significant relationship between SWC and δ^13^C of twig phloem WSOM than that between SWC and δ^13^C of leaf WSOM demonstrated that δ^13^C of twig phloem WSOM is a more sensitive indicator of SWC. Twig phloem WSOM was generally ^13^C-depleted compared with leaf WSOM for potential post-photosynthetic fractionation, and the ^13^C discrimination from leaves to twig phloem was insensitive to the interaction between SWC and C_a_. Clearly, interacting effects play a more important role in photosynthetic fractionation than in post-photosynthetic fractionation.

**Highlight:** The δ^13^C of leaf and twig phloem WSOM exhibited the most positive values at C_400_×35%–45% FC.

Post-photosynthetic fractionation from leaf to twig was not be impacted by the interacting effects.

## Introduction

Over the past 160 years, atmospheric CO_2_ (C_a_) concentration have increased from 280 μmol mol^-1^ to almost 400 μmol mol^-1^ and could surpass 700 μmol mol^-1^ with an annual rate of increase of 0.4% (Zhao *et al*., 2017). Concurrently, soil water stress may be aggravated by a climate that is both warming and drying, especially in regions where soil evaporation far exceeds precipitation. Within this context, it is important to investigate the interaction between soil water content (SWC) and C_a_ on plant performance and to predict plant fragmentation mechanisms under elevated C_a_ and heightened drought conditions. Carbon isotopes from the fast-turnover carbohydrate pool (water-soluble organic material, WSOM) in leaves are commonly utilized as an indicator of the assimilation-weighted average ratio of C_i_ (foliar intercellular CO_2_ concentration) to C_a_ carboxylated over a period of time, from several hours to 1–2 days (Pons *et al*., 2009). In addition, the leaf exported carbon in phloem carries the ^13^C signal of its source tissue. Therefore, the determination of carbon isotope composition (δ^13^C) in a plant is a promising tool that is commonly employed as an indicator of the effects of environmental factors on plant performance.

_δ_^13^C of WSOM is sensitive to subtle changes in environmental factors, including SWC and C_a_, as a result of variation in photosynthetic and physiological traits are imprinted in δ^13^C of new assimilates (Robredo *et al*., 2010; Adiredjo *et al*., 2014; Luo *et al*., 2016). However, the mechanism of how changes in environmental factors influence δ^13^C of twig phloem WSOM might be more complexity compared with that of leaf WSOM because δ^13^C of leaf-exported carbon in phloem can be altered by post-photosynthetic carbon isotope fractionation (Brandes *et al*., 2006; Kodama *et al*., 2011). Further, photosynthetic discrimination cannot completely explain the δ^13^C variation in different plant organs (Damesin and Lelarge, 2003; Bowling *et al*., 2008). For example, Gessler *et al*. (2008) reported that 30% of the differences in δ^13^C between leaf carbon and transported carbon can be explained by post-photosynthetic carbon isotope fractionation. In contrast to photosynthetic carbon isotope fractionation, post-photosynthetic carbon isotope fractionation has not been well defined or understood. Based on previous research, post-photosynthetic carbon isotope fractionation has been associated with activities such as transport of metabolites, the process of respiration, and biosynthesis of organic materials (Damesin and Lelarge, 2003; Gessler *et al*., 2004; Brandes *et al*., 2006; Bathellier *et al*. 2008).

The individual influence of SWC and C_a_ on δ^13^C of leaf and twig phloem WSOM has been under investigation; however, scant research has explored how δ^13^C of leaf and twig phloem WSOM responds to the interaction of SWC and C_a_. Assessing variation in δ^13^C of new assimilates in leaves and comparing δ^13^C of leaf exported organic materials in the twig phloem are effective methods for describing both photosynthetic and post-photosynthetic carbon isotope fractionation, providing valuable insights into plant physiology and biogeochemistry (Badeck *et al*., 2005). In this study, the interacting effects of SWC and C_a_ on δ^13^C of water-soluble leaf and twig phloem material were investigated by orthogonal treatments of three C_a_ levels coupled with five SWC gradients (three C_a_× five SWC). The objectives of this study were to describe the interaction between SWC and C_a_ on δ^13^C of water-soluble leaf and twig phloem material, and to determine whether ^13^C discrimination during the leaf-to-stem transition of carbohydrates was associated with the interacting effects of SWC and C_a_.

## Material and methods

### Experimental design

Research was conducted at the Chinese Research Station of Forest Ecosystems in the capital circle, located in Jiufeng National Forest Park of northwest Beijing (116°05´E, N40°03´N). In May 2016, 45 local three-year-old *Platycladus orientalis* saplings were selected for transplantation into pots (one sapling per pot). The top diameter of each pot was 30 cm, base diameter was 22 cm, and height was 29 cm. Soil with a field capacity (FC) of 26.2% was collected from the local undisturbed forest. The saplings had similar growth status, tree height (about 1.4 m), and root collar (about 28 mm). After two months of acclimation, the potted saplings were moved to growth chambers (FH-230, Taiwan Hipoint Corporation, Kaohsing City, Taiwan) for orthogonal treatments.

The design scheme of the orthogonal treatments (three C_a_ levels coupled with five SWC gradients) is presented in Table 1. There were two chambers used in the study for orthogonal treatments. One chamber was connected to a CO_2_ tank (Fig.1a), which can maintain C_a_ of 600 μmol mol^-1^ (C_600_) and 800 μmol mol^-1^ (C_800_). Another chamber was connected to an N_2_ tank (Fig.1b), which was used to maintain an ambient atmospheric concentration of approximately 400 μmol mol^-1^ (C_400_). The central controlling system of the chambers (FH-230) can consistently monitor and adjust C_a_ to their set values. Each chamber held five potted saplings and all five had a different SWC: the first was 35%–45% of FC (SWC between 9.17% and 11.79%), the second was 50%–60% of FC (SWC between 13.10% and 15.72%), the third was 60%–70% of FC (SWC between 15.72% and 18.34%), the fourth was 70%–80% of FC (SWC between 18.34% and 20.96%), and the fifth was 95%–100% of FC (SWC between 24.89% and 26.20%). Five saplings were cultivated at each SWC condition at each C_a_. Saplings in each treatment were allowed to acclimate for two weeks, and each treatment was repeated three times. To recreate local field conditions, the temperature and relative humidity in the growth chambers were set as follows: from 07:00 to 19:00, the lighting system was activated (photosynthetic photon flux density ranged between 200 and 240μmol m^-2^ s^-1^), temperature was 25±0.5 °C, and relative humidity was 60%. From 19:00 to 9:00, all light was turned off, temperature was 18±0.5 °C, and relative humidity was 80%.

**Figure 1.**
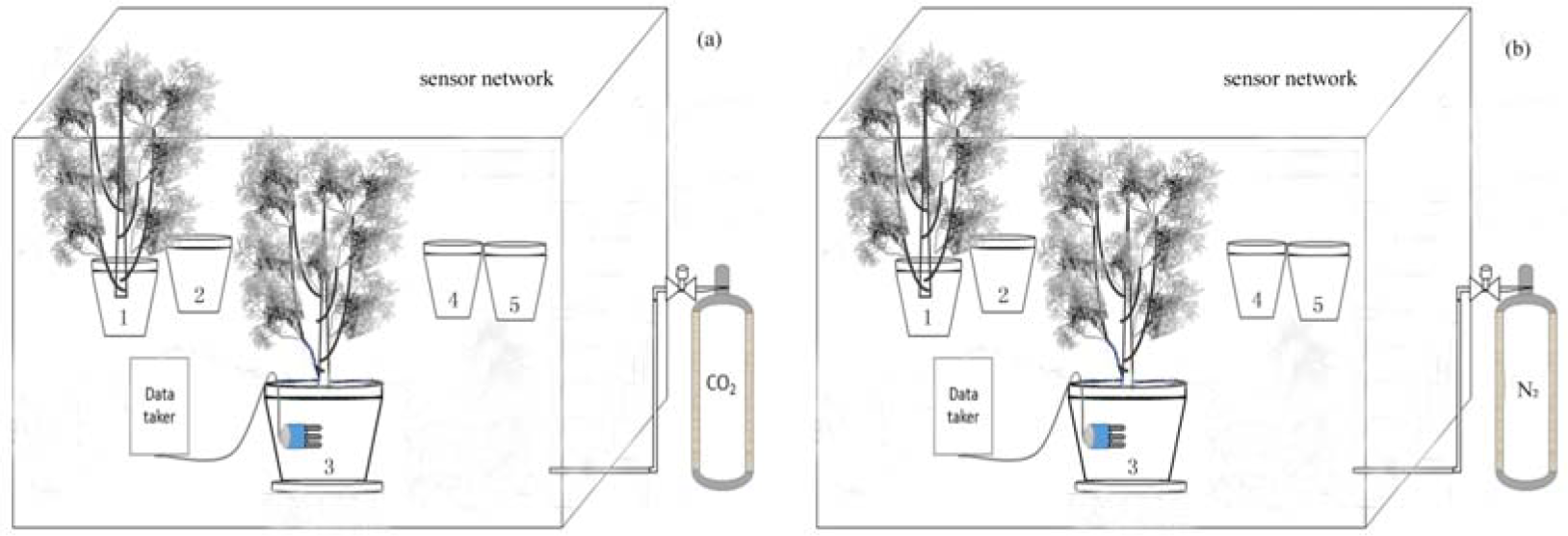
Growth chambers used in the experiments. One growth chamber (a) was used to maintain C_a_ of 600 μmol mol^-1^ and 800 μmol mol^-1^, another growth chamber (b) was used to maintain C_a_ of 400 μmol mol^-1^.

**Table 1.**
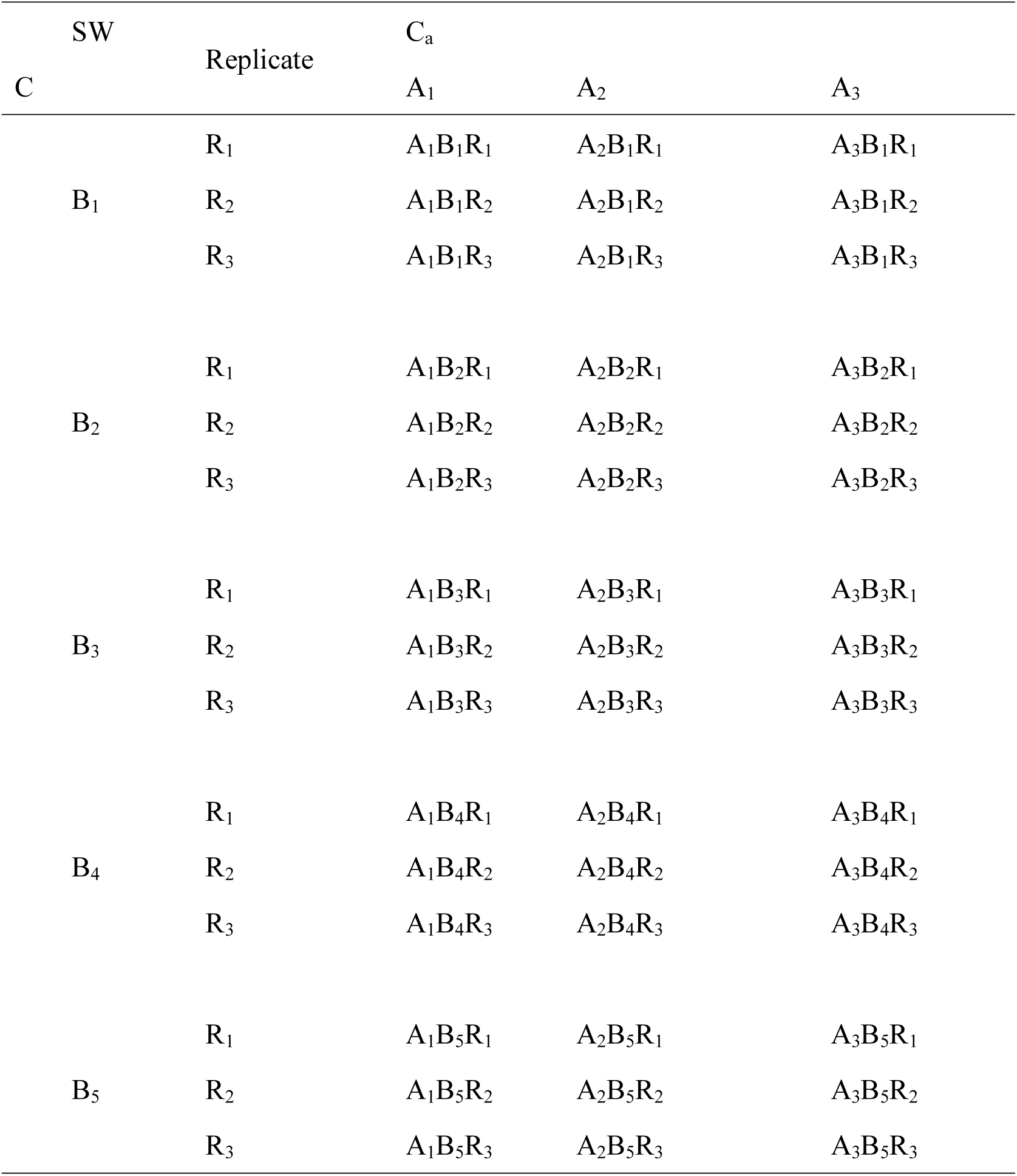
Scheme of the orthogonal experiment used in this research. A represents C_a_, and B represents SWC. A_1_, A_2_, and A_3_ are C_400_, C_600_, and C_800_, respectively. And B_1_, B_2_, B_3_, B_4_ and B_5_ are different SWC levels (35%–45% FC, 50%–60% FC, 60%–70% FC, 70%–80% FC, and 95%–100% FC, respectively). R represents the number of replicates of each treatment.

### Sampling and measurement

#### Foliar gas exchange measurement

Between 9:00 and 11:00, photosynthetic and physiological traits, including net photosynthetic rate (P_n_), stomatal conductance (g_s_), and foliar intercellular CO_2_ concentration (C_i_), were measured on fully expanded leaves using a portable photosynthesis system (Li-6400, Li-Cor, Lincoln, NE, US) after two weeks of treatment. Measurements were made on three leaves per sapling. Light conditions in the growth chamber remained stable, almost all leaves were exposed to a similar light intensity, and there was no obvious intracanopy gradient; therefore, the measurement of photosynthetic and physiological traits was only conducted in the upper parts of the saplings.

#### Sampling and δ^13^C of WSOM measurement

Following the measurements of gas exchange, the three measured leaves and three small twigs attached the leaves per sapling were collected, wrapped in tinfoil, and preserved in liquid nitrogen to prevent physiological activities.

To obtain leaf WSOM, 50 mg of freshly crushed leaf samples were incubated in 1.75 mL double demineralized water in a 2.0 mL centrifuge tube for one h at a constant temperature of 5 °C. Subsequently, the leaf material was boiled at 100 °C for 3 min to precipitate proteins and centrifuged for 5 min (12000 g) to obtain the supernatants for isotope analysis.

The twig phloem exudates were collected according to the method proposed by Gessler *et al*. (2004). Similar to the collection of leaf WSOM, 75 mg twig phloem samples were soaked in 1.75 mL double demineralized water in a 2.0 mL centrifuge tube at room temperature for 24 h. According to Schneider *et al*. (1996) and Bogelein *et al*. (2012), contamination of cellular constituents dissolving into the solution is negligible during this time period. The exudates were then centrifuged for 5 min (12000 g) to obtain supernatants.

The resulting supernatants of both leaf and twig material (10 μL) were placed in tin capsules and oven dried at 70 °C. These capsules were transported to the Institute of Botany of the Chinese Academy of Sciences for analysis. δ^13^C in water-soluble leaf and phloem material was analyzed using a stable isotope ratio mass spectrometer (DELTA^plus^XP, Thermo Finnigan), with a measurement precision of ±0.1‰.

On the plant sampling day, the atmosphere in the growth chambers was sampled using airbags and an air pump before harvest (at least three replicates). These airbags were also transported to the Institute of Botany of the Chinese Academy of Sciences for analysis. δ^13^C of atmospheric CO_2_ was analysed using a stable isotope ratio mass spectrometer (DELTA^plus^XP, Thermo Finnigan), with a measurement precision of±0.1‰.

### Carbon isotope theory and calculation of mesophyll conductance

Carbon isotopic composition is generally expressed as an isotope ratio where delta (δ) is defined as:

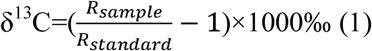

where R_standard_ and R_sample_ are the ^12^C/^13^C ratios of the standard and the sample.

It is well known that ^13^C discrimination occurs during the photosynthetic process, which results in different δ^13^C values between the atmosphere and the plant. The measured photosynthetic ^13^C discrimination between ambient air and leaf/twig is expressed as follows (Farquhar and Richards, 1984):

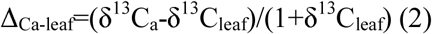

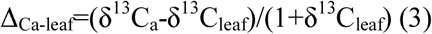

where δ^13^C_a_, δ^13^C_leaf_, and δ^13^C_twig_ are δ^13^C in ambient air, leaf WSOM, and twig WSOM, respectively.

In the most commonly used version of the Farquhar *et al*. (1982) simple model, the photosynthetic ^13^C discrimination was linearly related to C_i_/C_a_. Thus, photosynthetic ^13^C discrimination derived from gas exchange measurements (Δ_i_) is calculated as:

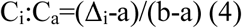

where a is the fractionation associated with CO_2_ diffusion in air (4.4‰; Seibt *et al*., 2008), and b is the fractionation relevant to reactions by Rubisco and PEP carboxylase (27‰; Hu *et al*., 2010).

Indeed, CO_2_ diffusion during photosynthesis includes three main steps: atmospheric CO_2_ reaches the surface of the leaf, it then enters into sub-stomatic cavities through the stomata, and it finally arrives at the chloroplast stoma in which photosynthesis occurs (Flexas *et al*., 2008). Therefore, photosynthetic ^13^C discrimination is dependent on the chloroplastic CO_2_ concentration (C_c_), rather than C_i_. Recently, studies have demonstrated that mesophyll resistance to CO_2_ diffusion would cause a drawdown from C_i_ to C_c,_ which means mesophyll conductance (g_m_) should be carefully considered (Flexas *et al*., 2006; Warren and Adams, 2006; Bogelein *et al*., 2012). The more comprehensive equation can be expressed as (Farquhar *et al*., 1982; Seibt *et al*., 2008):

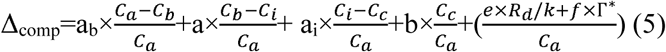

where C_b_ is the CO_2_ concentration on the leaf surface; a_b_ is the fractionation associated with atmospheric CO_2_ diffusion at the boundary layer (2.9‰); a_i_ is the fractionation for CO_2_ diffusion and dissolution in the liquid phase (1.8‰); e and f are the discrimination values for mitochondrial respiration (dark respiration, R_d_) and photorespiration, respectively; Г^*^ is the CO_2_ compensation point; and k is the carboxylation efficiency. It is generally accepted that the fractionation caused by boundary layer diffusion can be ignored and the equation can be simplified as (Shi *et al*., 2010):

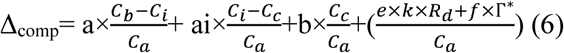

Substituting Δ_i_ into equation 6 we obtain the following equation:

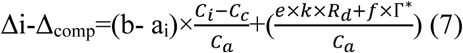

According to Fick’s first law, P_n_=g_s_×(C_a_-C_i_)=g_m_×(C_i_-C_c_) (Seibt *et al*., 2008). To calculate g_m_, we substituted Fick’s first law into equation (6) and the equation was transformed into:

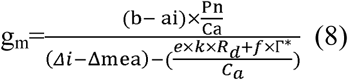

Ignorance of dark respiration and photorespiration can avoid further uncertainties and complexity, and including the two processes will not significantly improve model outcomes. For this reason, equation (8) can be simplified as:

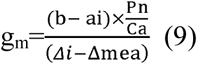

## Results

### Gas exchange properties

The gas exchange properties tended to increase with an increase in SWC, reaching a maximum at 70%–80% FC (Fig. 2). Under any given C_a_, the most striking increase in these gas exchange properties generally occurred when SWC increased from 35%–45% FC to 50%–60% FC. However, when SWC peaked at 95%–100% FC, these gas exchange properties began to decrease slightly (Fig. 2).

**Figure 2.**
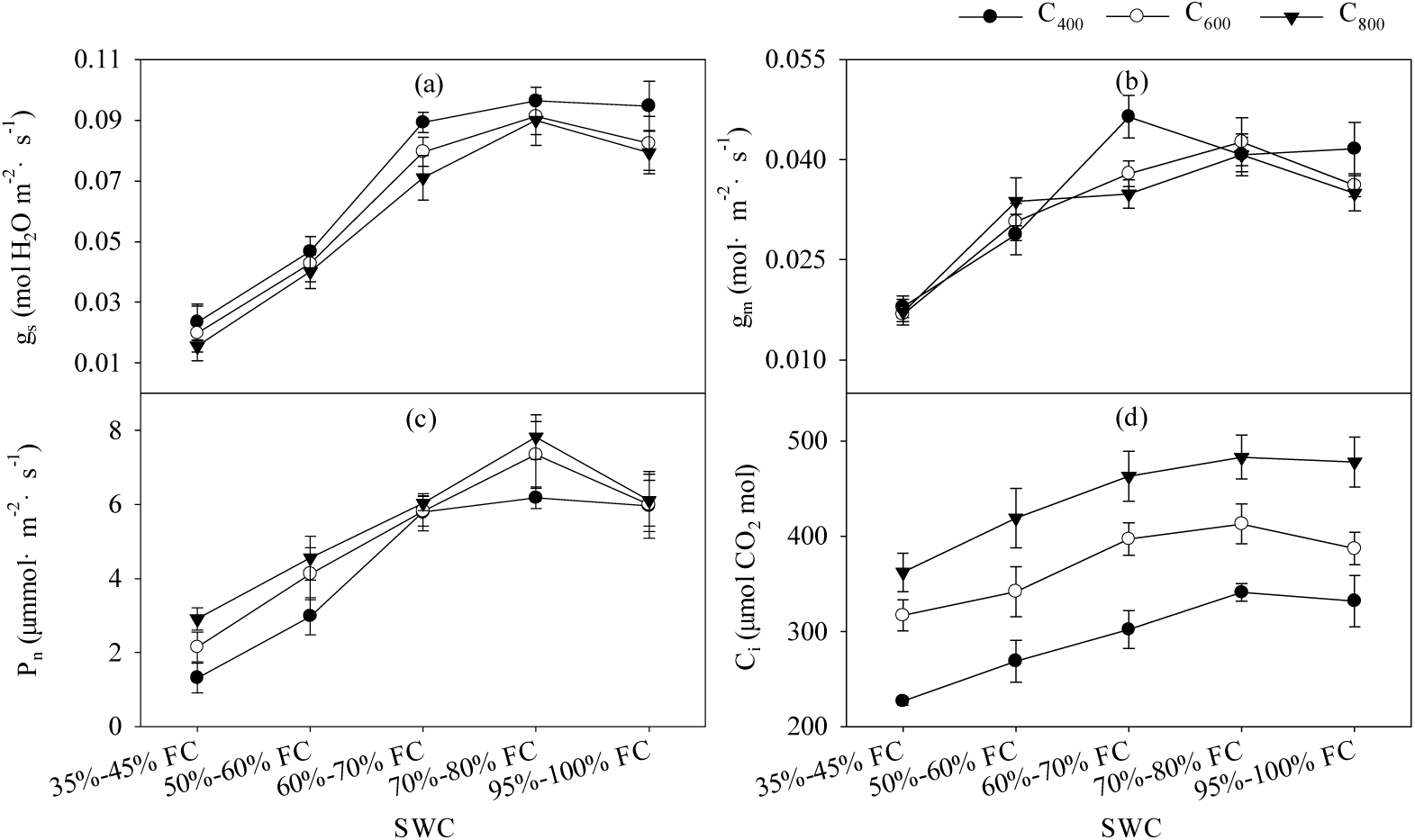
Gas exchange properties of *Platycladus orientalis* saplings under five SWC × three C_a_. Data are mean values ± SD.

Elevation of C_a_ caused an increase in P_n_ and C_i_ under any given soil moisture condition (Fig. 2c and 2d), while g_s_ decreased with the elevation of C_a_ (Fig. 2a). Furthermore, g_m_ tended to decline as C_a_ increased except when SWC was 50%–60%, which was similar to the pattern of change of g_s_ (Fig. 2b).

Correlation analysis revealed that P_n_ was significantly correlated with g_s_, g_m_^,^ and C_i_, with correlation coefficients (*r*) of 0.91, 0.90, and 0.72 (*p*<0.01), respectively (Table 2). In contrast, the relationship between P_n_ and C_i_/C_a_ was not strong or even moderate. Moreover, C_i_ and gs or gm were not significantly correlated, but both g_s_ and g_m_ were highly correlated with C_i_/C_a_ (*r* of 0.78 and 0.72, respectively; *p*<0.01). Additionally, g_m_ was closely correlated with g_s_ (*r* = 0.94; *p*<0.01) (Table 2).

**Table 2.** Correlation between gas exchange properties. n=15; * *p*<0.05, ** *p*<0.01.

| | $g_s$ | $g_m$ | $C_i$ | $C_i/ C_a$ | $P_n$ |
| --- | --- | --- | --- | --- | --- |
| $g_s$ | — | 0.94** | 0.42 | 0.78** | 0.91** |
| $g_m$ | | — | 0.41 | 0.72** | 0.90** |
| $C_i$ | | | — | — | 0.72** |
| $C_i/ C_a$ | | | | — | 0.49 |
| $P_n$ | | | | | — |

### δ^13^C of water-soluble leaf and twig phloem materials

_δ_^13^C of leaf WSOM ranged between -28.18‰ and -25.18‰, while δ^13^C of twig phloem WSOM had a wider range of -28.79‰ and -23.78‰ (Fig. 3). The δ^13^C of both leaf and twig phloem WSOM was maximized under the lowest SWC (35%–45% FC) at any given C_a_. When SWC increased from 35%–45% FC to 50%–60% FC, δ^13^C of water-soluble leaf and twig phloem materials had the most rapid decrease. Reduction of δ^13^C of water-soluble leaf and twig phloem materials typically leveled off when SWC was higher than 60%–70% FC. Under non-severe water stress conditions, δ^13^C of both leaf and twig phloem WSOM were usually highest when C_a_ was 600 μmol mol^-1^ (Fig. 3). In contrast, elevation of C_a_ produced increases in δ^13^C of water-soluble leaf and twig phloem materials under the lowest SWC (35%–45% FC).

**Figure 3.**
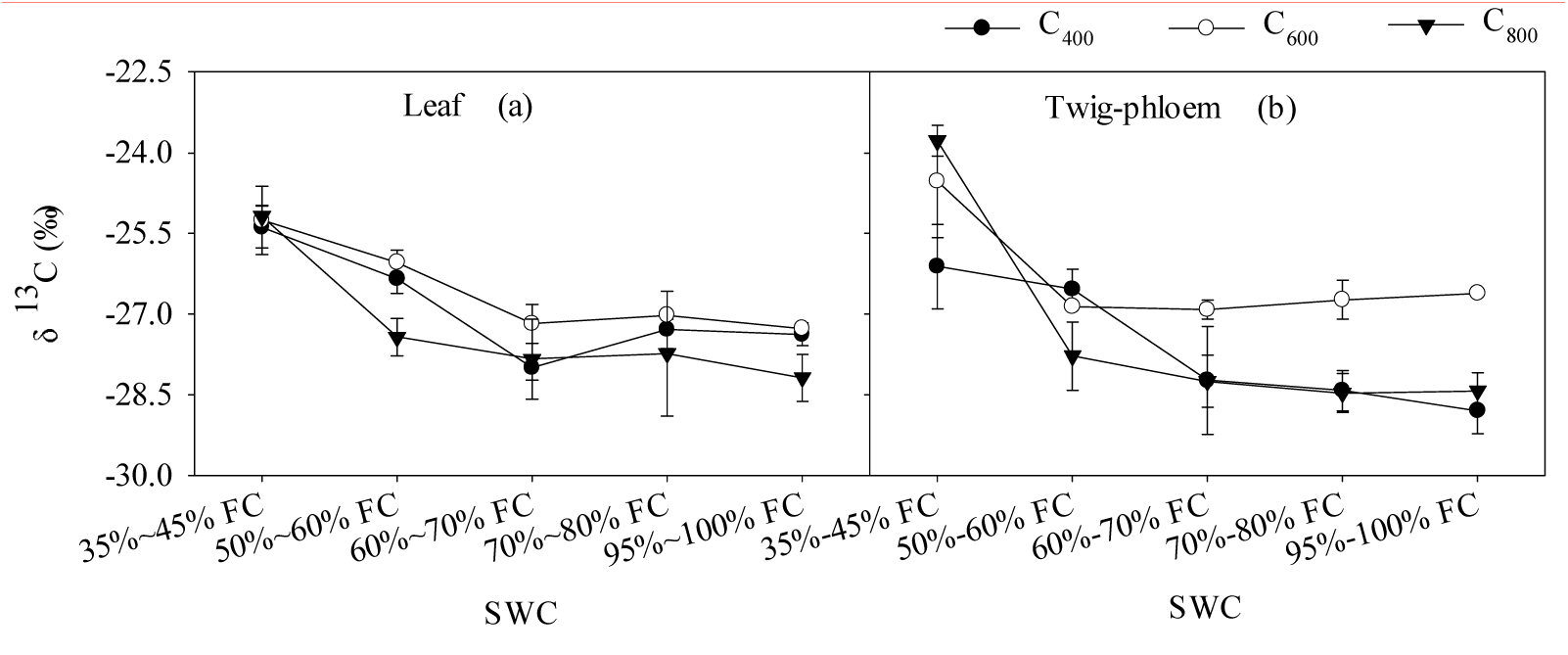
δ^13^C of water-soluble leaf (a) and twig phloem material (b) of *Platycladusorientalis* saplings under five SWC × three C_a_. Data are mean values ± SD.

### ^13^C enrichment from ambient air to plant

Δ_Ca-leaf_ ranged between 8.63‰ and 14.76‰, reaching a maximum at 60%–70% FC × C_400_ and a minimum at 35%–45% FC × C_800_ (Fig. 4a). In contrast, Δ_Ca-twig_ had a greater range between 7.18‰ and 15.59‰, reaching a maximum at 95%–100% FC × C_400_ and a minimum at 35%–45% FC × C_800_ (Fig. 4b). Under any C_a_, both Δ_Ca-leaf_ and Δ_Ca-twig_ tended to increase with increased SWC. Elevated C_a_ had an important influence on both Δ_Ca-leaf_ and Δ_Ca-twig_; Δ_Ca-leaf_ and Δ_Ca-twig_ decreased significantly when C_a_ increased from 400 μmol mol^-1^ to 600 μmol mol^-1^ (*p*<0.01). However, there were no significant differences in Δ_Ca-leaf_ and Δ_Ca-twig_ when C_a_ increased from 600 μmol mol^-1^ to 800 μmol mol^-1^ (*p*>0.05) (Fig. 4).

**Figure 4.**
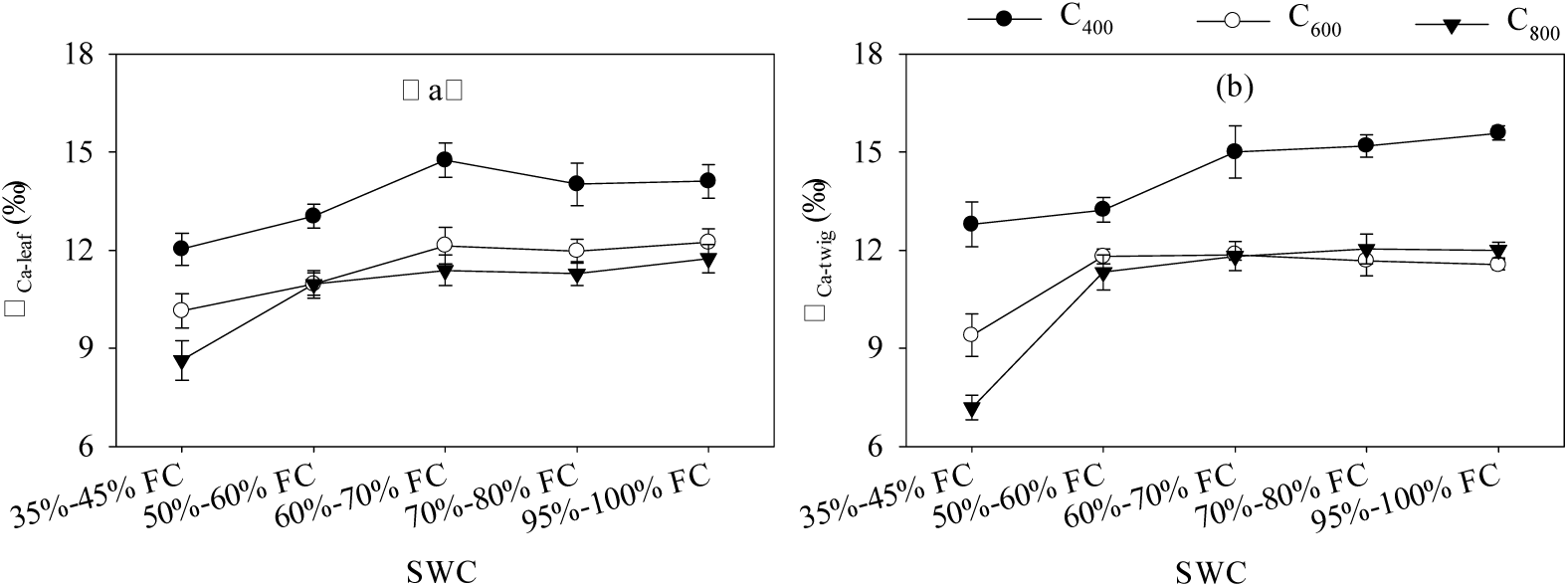
^13^C enrichment from ambient air to leaf (a) and twig (b) of *Platycladus orientalis* saplings under five SWC × three C_a_. Data are mean values ± SD.

### Correlation analyses

Analyses of correlation between gas exchange properties and δ^13^C and isotope discrimination are presented in Table 3. There were significant correlations between g_s_ and g_m_ and δ^13^C of leaf WSOM (*r* of -0.84 and -0.87, respectively; *p*<0.01). The δ^13^C of twig phloem WSOM was also highly correlated with g_s_ and g_m_ (*r* of -0.78 and -0.82, respectively; *p*<0.01). Moreover, similar significant correlations between g_s_ and g_m_ and Δ_Ca-leaf_ (for both, *r*=0.69, *p*<0.01) as well as between g_s_ and g_m_ and Δ_Ca-twig_ (*r* of 0.66 and 0.67, respectively; *p*<0.01) were observed. Table 3 also shows that C_i_ was moderately associated with δ^13^C of leaf WSOM (*r*=-0.62, *p*<0.05) but there was no significant or even moderate correlation between C_i_ and δ^13^C of twig phloem WSOM (*r*=-0.38, *p*>0.05). However, δ^13^C of twig phloem WSOM was highly related with C_i_/ C_a_ (*r*=-0.62, *p*<0.05). Additionally, C_i_/ C_a_ was strongly associated with Δ_Ca-leaf_ and Δ_Ca-twig_ (*r* of -0.92 and -0.88, respectively; *p*<0.01).

**Table 3.** Correlation between gas exchange properties and δ13 538 C and carbon isotope discrimination. n=15; * *p*<0.05, ** *p*<0.01.

| | $\delta^{13}\text{C}_{\text{leaf}}$ | $\delta^{13}\text{C}_{\text{twig}}$ | $\Delta_{\text{Ca-leaf}}$ | $\Delta_{\text{Ca-twig}}$ |
| --- | --- | --- | --- | --- |
| $g_s$ | -0.84** | -0.78** | 0.69** | 0.66** |
| $g_m$ | -0.87** | -0.82** | 0.69** | 0.67** |
| $C_i$ | -0.62* | -0.38 | — | — |
| $C_i/ C_a$ | -0.48 | -0.62* | -0.92** | -0.88** |

## Discussion

### ^13^C fractionation from leaves to twig phloem

We observed no clear patterns of variation in ^13^C fractionation from leaves to twig phloem in response to the interaction between SWC and C_a_. In most cases, ^13^C was depleted in the twig phloem WSOM in comparison to leaf WSOM (Fig.5), indicating that δ^13^C of fast turnover carbon can be altered as a result of post-photosynthetic fractionation. Such depletion was also reported by Gessler *et al*. (2004) for beech. However, Bogelein *et al*. (2012) found that there was ^13^C enrichment in twig phloem exudates in comparison to leaf WSOM in the upper and middle beech canopy, but such enrichment was not detected in the lower shaded beech canopy. Similar intra-canopy patterns of twig-to-leaf δ^13^C differences were also reported by Cernusak *et al*. (2009). Further, Eglin *et al*. (2009) observed that WSOM had higher δ^13^C values in twigs than in leaves within sunlit beech crowns.

**Figure 5.**
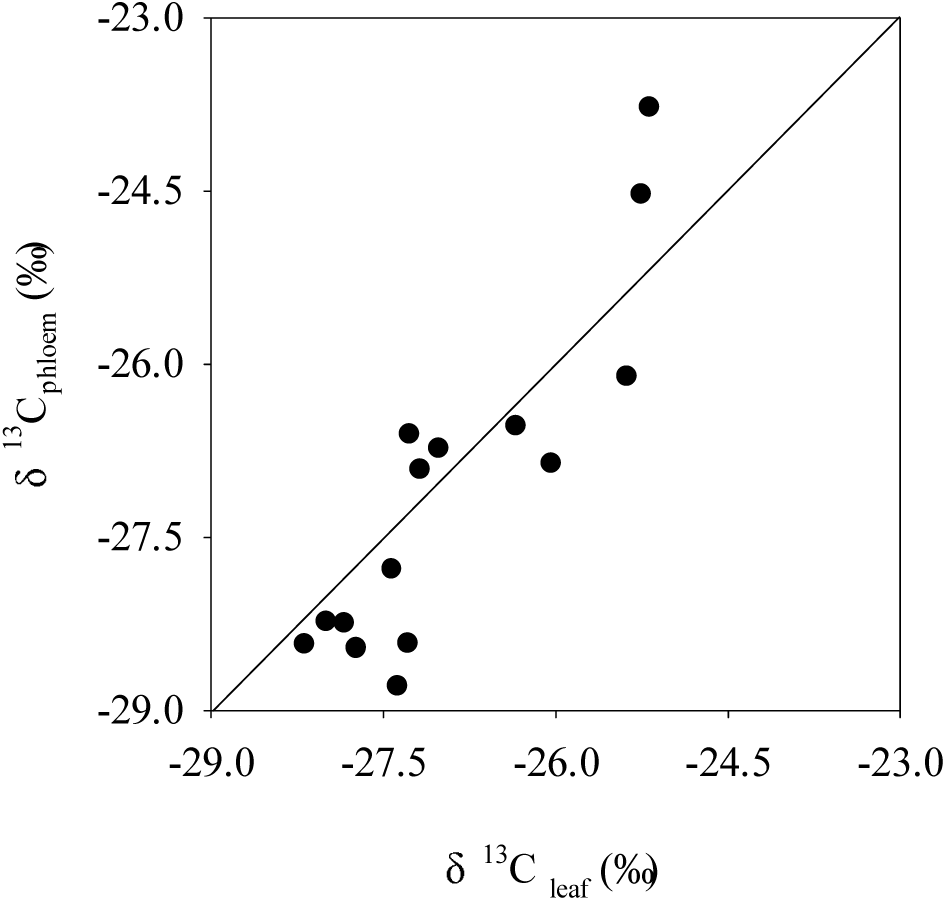
The δ^13^C of leaf WSOM versus δ^13^C of twig phloem WSOM.

It is well known that a large amount of glucose is synthesized in chloroplasts when photosynthesis is active, part of which is exported to the cytosol by triose phosphate translocators (TPTs) for sucrose biosynthesis and other metabolic activities; excess glucose accumulates and is converted into starch in chloroplasts (Cho *et al*., 2011; Mahboubi, 2014). During this process, the aldolase isotope effect leads to relatively ^13^C enriched hexoses being incorporated in starch while isotopically lighter trioses will be exported to the cytoplasm. Therefore, δ^13^C in the chloroplastic starch was more positive than that in the cytosolic WSOM for plants with strong photosynthetic capacity (Brandes *et al*., 2006; Cernusak *et al*., 2009). In contrast, the remobilization of starch in chloroplasts as a result of lower photosynthetic rates leads to sucrose exported from leaves carrying the signature of the starch substrate. Previous studies determined that sucrose is the major carbohydrate that can be exported into phloem and transported throughout the whole plant (Gessler *et al*., 2008; Mahboubi, 2014). Indeed, Peuke *et al*. (2001) observed that about 90% of the carbohydrates transported in the phloem consists of sucrose. This might result in differences in sunlit versus shaded crowns in ^13^C fractionation patterns of leaves to twig phloem. In this study, the daytime light intensity in the growth chambers ranged from 200 and 240μmol m^-2^ s^-1^, which is a much smaller range than that of field conditions (ranging from 40 to 1500μmol m^-2^ s^-1^ with an average of about 350μmol m^-2^ s^-1^). Therefore, the δ^13^C difference between leaf and twig phloem WSOM in this study could be because of the absence of starch storage under low light intensity in the growth chambers.

However, the potential post-photosynthetic carbon isotope fractionation mechanism can be species-specific. For example, Bogelein *et al*. (2012) claimed that twig phloem WSOM was ^13^C enriched compared with leaf WSOM throughout the whole canopy for Douglas fir. In a review of more than 80 studies, Badeck *et al*. (2005) found that autotrophic tissues were generally ^13^C-depleted compared with heterotrophic tissues. According to Merchant *et al*. (2010) and Cernusak *et al*. (2009), different proportions of apoplastic and symplastic loading pathways could independently influence δ^13^C, introducing a potential species-independent factor in the fractionation of phloem isotope composition in contrast to the source tissue. In addition, twig phloem WSOM is a mixture of carbohydrates with different metabolic histories derived from carbon exported from the source tissue (leaves) during both light and dark periods. Cernusak *et al*. (2009) postulated that if leaves and small twigs were sampled at the same time during the day, twig phloem could carry ^13^C signals originating from both day and night exports from source tissues. Specifically, Ruehr *et al*. (2009) suggested that carbohydrate exportation takes less than 5 hours when SWC is high but up to 4 days under drought conditions. This may cause the ^13^C fractionation pattern from leaf to phloem to be more complicated. Previous studies also proposed that δ^13^C differences between leaf and twig phloem WSOM could be associated with respiratory processes (Cernusak *et al*., 2009; Merchant *et al*., 2010). However, Eglin *et al*. (2009) observed that there were no significant differences in δ^13^C of respired CO_2_ between leaves and twigs. This finding indicated that divergence in δ^13^C between the two organs is not likely to be explained by respiration fractionation. We suggest that post-photosynthesis fractionation may not be attributed to a single, unifying hypothesis; instead, it is the result of multiple processes. A better description of the characteristics of these hypotheses and what percentage of each processes is responsible for post-photosynthesis fractionation in different plant organs will improve our understanding of metabolism and plant physiology.

### Variations in δ^13^C of leaf and twig phloem WSOM

Variation in δ^13^C of leaf WSOM has been widely used to reveal plant leaf-level responses to environmental conditions (Hasselquist *et al*. 2010; Tomas *et al*. 2012). In this study, the effect of the interaction between SWC and C_a_ on δ^13^C of leaf WSOM was analysed. At any C_a_, δ^13^C of leaf WSOM was maximized under the lowest SWC (35%–45% FC). The effect of drought on stable carbon isotope composition at the leaf level of *Platycladus orientalis* saplings was similar to the effect on tolerant and sensitive beech ecotypes (Peuke *et al*., 2006). In addition, Nie *et al*.(2014) found that leaf δ^13^C signals of nearly all selected species growing in subtropical China exhibited were more positive during the dry season than in the wet season. When SWC increased to more than 50% FC, δ^13^C of leaf WSOM was maximized at C_600_. The δ^13^C of leaf WSOM didn’t increase if C_a_ get too high. The interactive effect between SWC and C_a_ on δ^13^C of twig phloem WSOM is similar to that on δ^13^C of leaf WSOM. Further regression analysis showed that the correlation between SWC and δ^13^C of twig phloem WSOM is significantly stronger than that between SWC and δ^13^C of twig phloem WSOM (Fig. 6). This result indicates that δ^13^C of twig phloem WSOM is a better predictor of SWC than that of leaf WSOM. Similar results have also be found by Merchant *et al*. (2010), who suggested that the soluble leaf carbon, amino acid, and nutrient concentrations in the leaf had lower sensitivity for the prediction of plant water status than δ^13^C of twig phloem WSOM, although δ^13^C of leaf sugars and soluble extracts mirrored the variation in δ^13^C of twig phloem WSOM. A larger or smaller time lag for twig phloem carbon, which is a mixture of various carbohydrates with different metabolic histories, is insufficient to override the ^13^C signal of newly synthesized carbohydrates (Merchant *et al*., 2010). Further, the phloem loading pathways by which new assimilates are exported from the leaf could be influenced by environmental conditions (Turgeon *et al*., 2001). This finding results from the phloem loading pathways being associated with membrane potential of mesophyll cells (Turgeon and Medville, 2004), and the pressure is sensitive to variations in the concentrations of sucrose and raffinose (van Bel *et al*., 1996). Phloem sap is dominated by sucrose and raffinose (Tausz *et al*., 2008), both of which are associated with plant water status (Merchant *et al*. 2010). Therefore, variations in SWC may alter the contributions of apoplastic and symplastic loading pathways, which affect δ^13^C of twig phloem WSOM.

**Figure 6.**
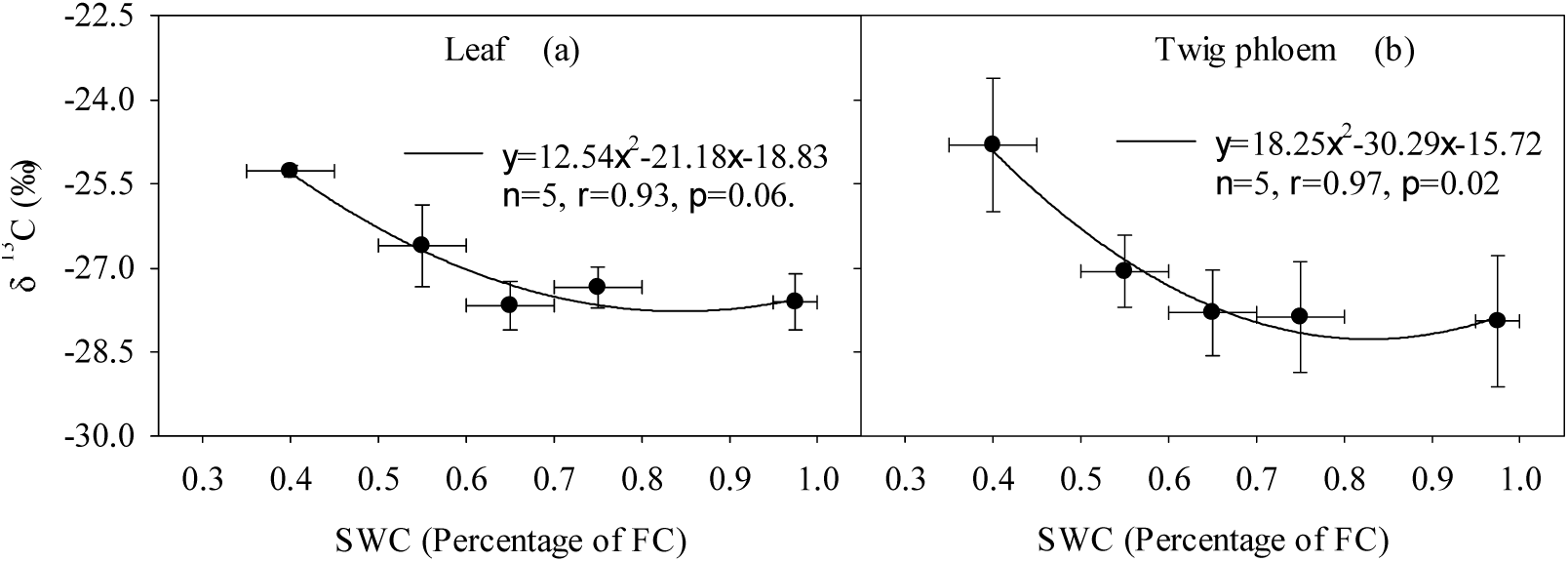
Relationship between SWC and δ^13^C of leaf and twig phloem WSOM.

### Correlations between carbon isotope discrimination and gas exchange properties

To date, numerous studies have confirmed the negative relationship between Δ_Ca-leaf_ and water use efficiency at the leaf level (Medrano *et al*., 2009; Cornejo-Oviedo *et al*., 2017), as Δ_Ca-leaf_ is highly correlated with C_i_/C_a_ (Farquhar *et al*., 1982; Table 3). We found that peak Δ_Ca-leaf_ was reached at 60%–70% FC × C_400_, and the minimum value was found at 35–45% FC × C_800_ (Fig. 4). This finding indicated more conservative water use strategies under higher C_a_ and drought conditions. Under elevated C_a_, the inhibition of photosynthetic uptake caused by drought can be alleviated, while transpiration capacity may remain constant or decrease (Bernacchi *et al*., 2006; Miranda Apodaca *et al*., 2015), which caused Δ_Ca-leaf_ to decrease. In addition, we found that Δ_Ca-twig_ is significantly correlated with C_i_/C_a_, suggesting that Δ_Ca-twig_ is a sensitive predictor of plant water use efficiency.

Carbon isotope discrimination from ambient air to the plant is dependent on g_s_ as C_i_ is regulated by g_s_, and simultaneously, g_s_ is subject to the feedback of carbon isotope discrimination. Early studies generally accepted that net photosynthesis depends on the supply of CO_2_ in the sub-stomata and ignored the important role that g_m_ played (Lauer *et al*., 1992). At that time, g_m_ was considered to be infinitely large, which resulted in C_i_ being equal or close to C_c_. However, more recent research has shown that g_m_ is a species-specific and environmentally dependent factor, and ignoring the potential difference between C_i_ and C_c_ could result in the erroneous interpretation of ^13^C discrimination. Therefore, caution must be exercised when considering the mechanism of ^13^C enrichment during photosynthetic CO_2_ fixation (Flexas *et al*., 2006; Seibt *et al*., 2008; Bogelein *et al*., 2012). Correlation analysis showed that g_s_ and g_m_ had similar positive effects on Δ_Ca-leaf_ and Δ_Ca-twig_ (Fig. 5). This finding emphasizes the important influence of g_m_ on ^13^C discrimination during photosynthesis. The similar influence that g_s_ and g_m_ had on Δ_Ca-leaf_ has also been reported by Shi *et al*. (2010). However, Warren (2004) claimed that mesophyll resistance had far more influence than stomatal resistance on CO_2_ diffusion during photosynthesis for *Eucalyptus globulus*, so it is possible that the mechanism might be species-specific.

## Acknowledgements

This study was supported by the National Natural Science Foundation of China (No.41430747), the National Science Fund for Distinguished Young Scholars (No.41401013), and the Beijing Municipal Education Commission (CEFF-PXM2017_014207_000043).

